# PhyloFusion- Fast and easy fusion of rooted phylogenetic trees into rooted phylogenetic networks

**DOI:** 10.1101/2024.06.25.600638

**Authors:** Louxin Zhang, Banu Cetinkaya, Daniel H. Huson

## Abstract

Unrooted phylogenetic networks are often used to represent evolutionary data when incompatibilities are present. Although rooted phylogenetic networks are better suited for explicitly depicting evolutionary histories that involve reticulate events, they have been rarely used in practice, due to a lack of appropriate methods for their calculation. Here we present PhyloFusion, a fast and easily-applicable method for calculating rooted phylogenetic networks on sets of rooted phylogenetic trees. The algorithm can handle trees with unresolved nodes (which arise when edges with low support are contracted) and missing taxa. We illustrate how to use the algorithm to explore different groups of functionally-related genes and report that the algorithm can be applied to datasets containing tens of trees and hundreds of taxa.

**Availability:** An open source implementation of PhyloFusion is available in SplitsTree, https://www.github.com/husonlab/splitstree6 (GPLv3 license)

## 1 Introduction

The evolutionary history of a set of species or other taxonomic groups is usually represented by a phylogenetic tree. However, when reticulate evolutionary events such as horizontal gene transfer, speciation by hybridization, recombination, reassortment or incomplete lineage sorting are suspected to play a role, then methods that compute a phylogenetic network may be more appropriate [Huson et al., 2012].

In the case of unrooted networks, Neighbor Net [Bryant and Moulton, 2004] and Median Joining [Bandelt et al., 1999] are widely used and cited, and are provided by the SplitsTree app [Huson and Bryant, 2024]. However, in the case of rooted networks, a general purpose, easily applicable tool has been lacking [Stolzer et al., 2012, Huson and Linz, 2018, van Iersel et al., 2022].

Recently, Zhang et al. [2023] introduced a new algorithm for computing a rooted phylogenetic network for an input set of rooted phylogenetic trees. The computed network has the *tree-child* property, which means that every internal node of the network has at least one child that is a tree node. Their ALTS algorithm can be applied to tens of input trees and computes a rooted network is seconds. Alas, their algorithm requires that all input trees are fully resolved. This poses a severe limitation in practice, because low-confidence edges in the input trees usually give rise to unnecessarily complicated and distracting reticulation scenarios in the output. Also, the algorithm is not designed to allow missing taxa in the input trees.

Here we introduce *PhyloFusion*, an extension of the ALTS algorithm. PhyloFusion is a fast and easily-applicable method for exploring the use of rooted phylogenetic networks in the context of gene trees (or other tree collections). Like ALTS, this algorithm takes as input a list of rooted phylogenetic trees and produces as output a rooted phylogenetic network that contains all the input trees, heuristically minimizing the hybridization number associated with the network. Improving on ALTS, the PhyloFusion algorithm is designed to accept unresolved nodes and missing taxa in the input trees. The algorithm is fast enough to allow tens of trees and hundreds of taxa as input and thus can be used to interactively explore sets of related gene trees.

To illustrate the use of PhyloFusion, we apply it to different sets of chloroplast genes trees for water lilies [Gruenstaeudl, 2019]. While a rooted phylogenetic network encompassing all 78 trees is perhaps too complicated to be useful, the networks computed on smaller sets of functionally related genes give rise to much simpler, potentially useful reticulation scenarios.

## 2 Results

We provide an implementation of the *PhyloFusion* algorithm in the SplitsTree app [Huson and Bryant, 2024]. Our algorithm takes as input a set of rooted phylogenetic trees and produces, as output, a rooted phylogenetic network that contains all input trees. The algorithm is a heuristic that aims at minimizing the hybridization number of the network.

Basic features of SplitsTree can be used to preprocess the data. First, the *Reroot or Reorder* method allow one to ensure that all trees are correctly rooted, either by outgroup or midpoint rooting. Second, the *Trees Filter* method can be used to select a subset of trees to place into the network. Third, the *Tree Edges Filter* method should be used to contract all edges that have low confidence (such has less than 70% bootstrap support, say). The *PhyloFusion* algorithm can then be run on the preprocessed trees. By default, our implementation of the algorithm will first perform *mutual refinement* of all input trees in an attempt to remove topological differences in the input trees that are only due to different degrees of resolution.

We envision researchers using PhyloFusion to explore the application of rooted phylogenetic networks on sets of functionally related genes. To illustrate this, we use 43 photosynthesis-related chloroplast gene trees for 17 waterlilies, based on multiple sequence alignments provided in [Gruenstaeudl, 2019]. For each of these alignments, we computed a maximum-likelihood tree with bootstrap values using IQ-TREE [Minh et al., 2020].

In a preprocessing step, all trees where rerooted using the two Trithuria species as outgroup and then all edges with a bootstrap support of less than 70% were contracted. Then, we ran PhyloFusion on all 43 photosynthesis-related gene trees and obtained a rooted phylogenetic network with hybridization number *h* = 13, shown in Figure 1A. In a more detail analysis, there are 5, 15, 6 and 11 genes associated with Photosystem I, Photosystem II, Cytochrome b6/f Complex, ATP Synthase and NDH Complex, respectively, and the corresponding rooted phylogenetic networks have hybridization number 3, 8, 4, 2 and 5, respectively, as shown in Figure 1B-F.

**Figure 1.**
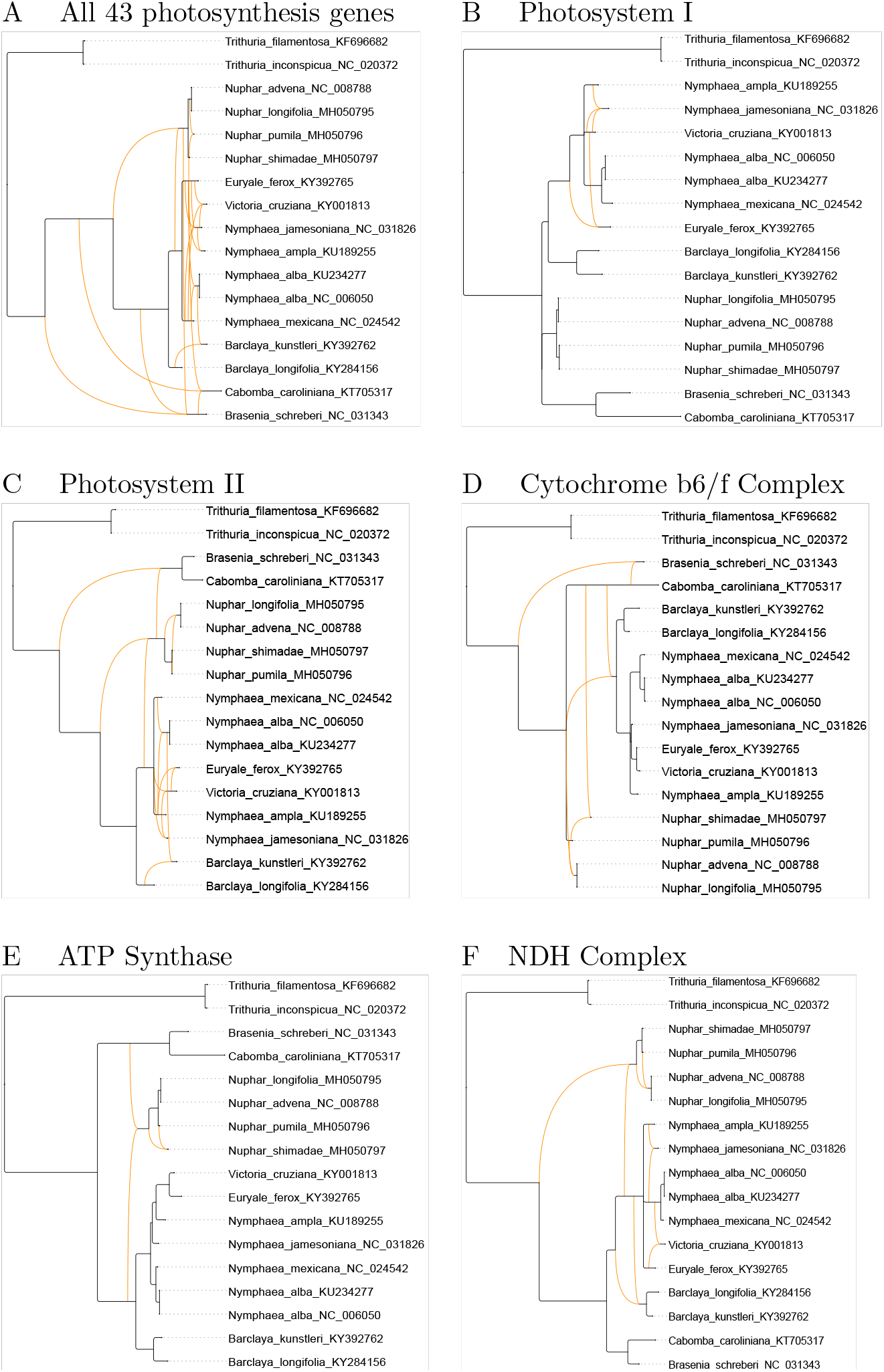
For rooted chloroplast gene trees on 17 taxa [Gruenstaeudl, 2019], using a minimum bootstrap threshold of 70% and the PhyloFusion algorithm, we show: A: a rooted phylogenetic network containining all 43 photosynthesis-related gene trees (hybridization number *h* = 13), B: containing all 5 *psa* (Photosystem I) gene trees (*h* = 3), C: containing all 15 *psb* (Photosystem II) gene trees (*h* = 8), D: containing all 6 *pet* (Cytochrome b6/f Complex) gene trees (*h* = 4), E: containing all 6 *atp* (ATP Synthase) gene trees (*h* = 2), and F: containing all 11 *ndh* (NDH Complex) gene trees (*h* = 5), respectively.ss

In each of the cases, the calculation of the network takes at most a few seconds. While we do not suggest that reticulate evolution is a key factor here, the dataset allows us to demonstrate how one may use PhyloFusion to explore the use of rooted phylogenetic networks to represent the evolutionary history of sets of related genes in a straight-forward manner. Running PhyloFusion on the full set of all 78 chloroplast gene trees takes less than a minute and processes a network with *h* = 31.

To systematically evaluate the performance of PhyloFusion, we ran the algorithm on datasets ranging from 10 to 100 taxa, with 2 to 10 input trees, each obtained by applying one random “rooted Subtree Prune and Regraft” (rSPR) operation to a fixed source tree. In Figure 2A, we plot the wall-clock time required. In Figure 2B, we compare the hybridization number of the computed networks to the total number of rooted SPRs contained in the input trees.

**Figure 2.**
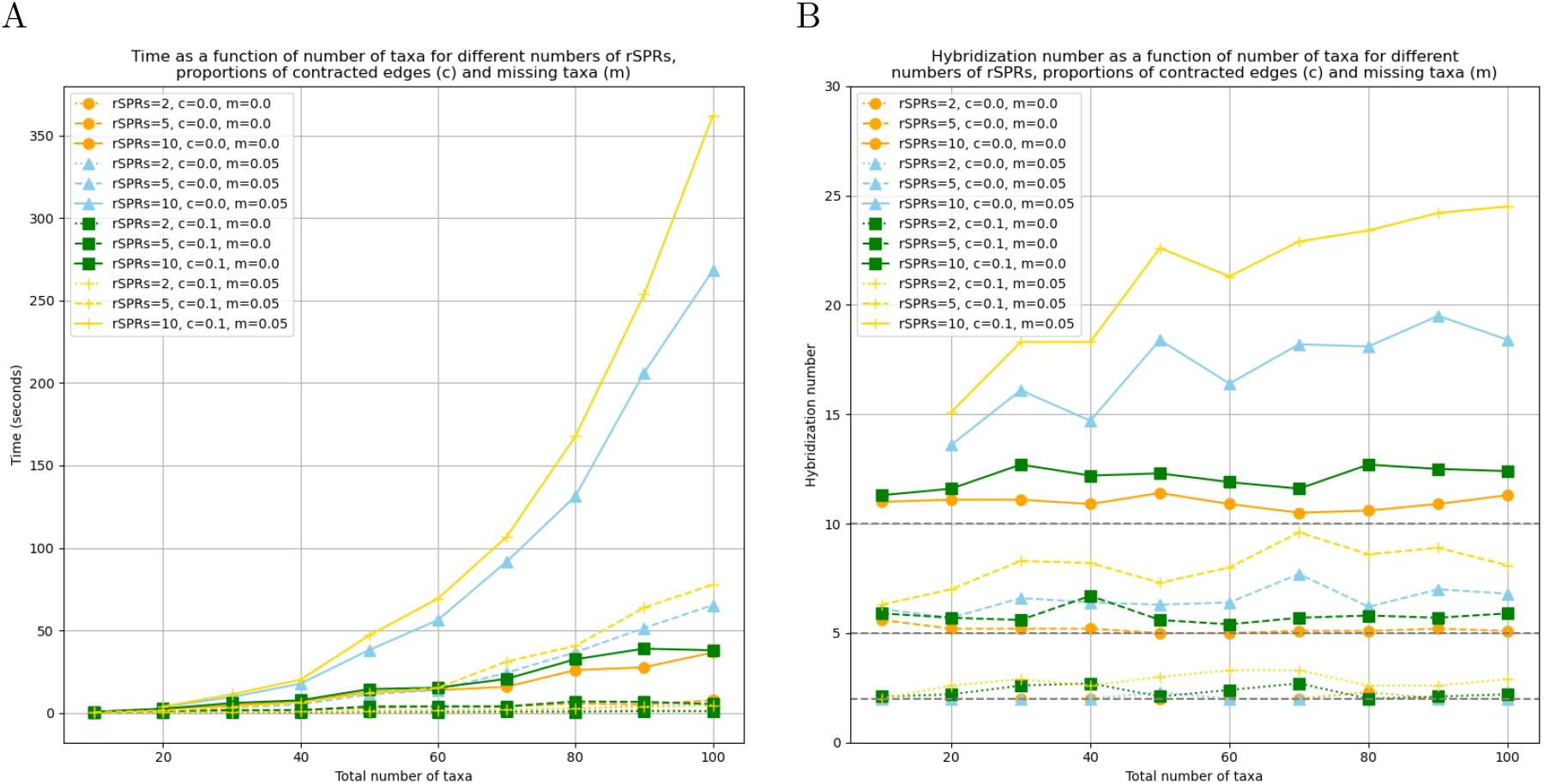
A: For 2, 5 and 10 input trees, each containing one rSPR, here we plot the number of seconds required by PhyloFusion, as a function of the number of taxa in each of the trees. B: Here we plot the hybridization number of the computed networks, as a function of the number of taxa in the trees. Each plotted point shows the average of 10 replicates. The numbers *c* and *m* indicate the proportion of contracted internal edges and missing taxa, in each input tree, respectively.

The plots suggest that the method performs well on realistic problem sizes. Note that simulating missing taxa and simulating unresolved nodes both increase the minimum hybridization number required to encompass all trees. In the case of missing taxa, consider, for example, three taxa *a, b, c* whose true rooted phylogeny is given by ((*a, b*), *c*). If each one of the three taxa is missing in one of the input trees, then the input dataset will contain all three sibling pairs (*a, b*), (*a, c*) and (*b, c*) and will therefore require a reticulation to encompass them. For larger numbers of trees and taxa, and a moderate proportion of missing taxa, such triplets are very common. In the case of unresolved nodes, note that the algorithm aims at producing a network that contains every input tree in the strict sense that it contains the tree itself, and not merely a resolution of it. In consequence, if one input tree contains an unresolved node that is resolved in an other input tree, then this will require a reticulation in the output network so as to accommodate both trees. In an optional preprocessing step, PhyloFusion attempts to reduce the number of reticulations due to this by performing mutual refinement of the input trees.

## Discussion

While unrooted phylogenetic networks are widely used in the literature, there are few examples of studies in which a rooted phylogenetic network is used to represent a putative reticulate evolutionary history, such as [Koblmueller et al., 2007]. One reason for this has been the lack of fast and easily-applicable methods for computing such networks. Here we describe PhyloFusion and show that it can be used to explore the application of rooted phylogenetic networks to collections of rooted phylogenetic trees, with the goal of fusing the trees into an encompassing network. Here we used a publish set of chloroplast genes [Gruenstaeudl, 2019] to demonstrate the usage of the algorithm on different collections of gene trees.

We envision PhyloFusion being used to explicitly represent cases of reticulate evolution. One potential area of application might be to eukaryotic datasets that have both organelle and nuclear gene trees, where cases of speciation by hybridization might be clearly visible in the calculated network. Another potential area of application might be to segmented RNA viruses, such as Influenza viruses, where reassortment may lead to significantly different topologies among trees based on different segments, which may be evident in a rooted network produced by PhyloFusion.

## 3 Methods

PhyloFusion is an extension of the ALTS algorithm [Zhang et al., 2023], which we first summarize. The input for ALTS is a set *T* = {*t*_1_, …, *t*_*k*_} of bifurcating phylogenetic trees, each containing the full taxon set *X* = {*x*_1_, …, *x*_*n*_}. The basic idea of the algorithm is that, for a given ordering of taxa, each pair of taxon *x* and tree *t*_*j*_ gives rises to a string *lts*(*x, t*_*j*_). For each taxon, these strings are merged into a shortest common super sequence and a network is then constructed from the merged sequences in such a way that the network is guaranteed to contain all trees.

In more detail, let *π* = (*x*_*π*(1)_, *x*_*π*(2)_, …, *x*_*π*(*n*)_) be a total ordering of *X*. The algorithm creates a phylogenetic network *N* = (*V, E*) with root node *ρ* and leaf labels *λ* : *V* → *X* as follows.

0 In a preprocessing step, ensure that all input trees are correctly rooted, for example using outgroup rooting. For algorithmic purposes, ensure that every input tree has a root node of out-degree 1 by inserting a new root above the original root, if necessary. Subtree reductions and clade reductions are applied recursively during processing [Zhang et al., 2023].
1 For each tree *t ∈ T* define a labeling *𝓁* of all nodes bottom-up as follows: If *u* is a leaf node, then set *𝓁*(*u*) = *x*, where *x* is the taxon associated with *u*, or, if *u* is the root, then set *𝓁*(*u*) = *x*_*π*(1)_. Otherwise, *u* is an internal node with two children, *v* and *w*. Let *m*(*v*) = min_*π*_ (*v*) and *m*(*w*) = min_*π*_ (*w*) be the smallest taxa associated with leaves below *v* and *w*, respectively. Then set *𝓁*(*u*) = max{*m*(*v*), *m*(*w*)}. This labeling has the property that each taxon *x* appears exactly twice, once as the label *𝓁*(*p*(*x*)) of an internal node *p*(*x*) and once as the label *𝓁*(*q*(*x*)) of a leaf node *q*(*x*), respectively [Zhang et al., 2023].
2 For each taxon *x*, produce a (single list of tax)a in two steps. First, for each tree *t*_*j*_ *∈ T*, create the sequence of taxa *lts*(*x, t*_*j*_) = *𝓁*(*v*_1_), …, *𝓁*(*v*_*k*_) that are associated with the nodes *v*_1_, …, *v*_*k*_ in the interior of the path from *p*(*x*) to *q*(*x*), this is called a “lineage taxon string” (LTS) in [Zhang et al., 2023]. Second, create a shortest common super sequence *scss*(*x*) that contains all the sequences *lts*(*x, t*_1_), …, *lts*(*x, t*_*k*_), obtained from the input trees for taxon *x*.
3 For each taxon *x*, let *L*(*x*) be the list of length |*scss*(*x*)| + 2 obtained by placing a copy of *x* both before and after the list *scss*(*x*). For each taxon *y ∈ L*(*x*), create a node *v* with label *𝓁*(*v*) = *y*. If *y* is the last element in *L*(*x*), then *v* will represent the taxon *x* and set *λ*(*v*) = *x*, else connect *v* to the next created node.
4 Construct a rooted phylogenetic network *N* as follows. The node *ρ* with label *𝓁*(*ρ*) = *x*_*π*(1)_ is declared the root of *N*. Then, for each taxon *x* in the order of *π* and for each internal node *v* in the path from *p*(*x*) to *q*(*x*), connect *v* to the first node *w* with *x*(*w*) = *𝓁*(*v*). For any node *v* that has precisely one parent and one child, create a new edge from the parent to the child, and delete *v*. This works because each taxon *x* (except the first one) is mentioned in the path associated with some other taxon *x*^*′*^ that appears before *x* in the taxon order *π* [Zhang et al., 2023].

The ALTS algorithm aims at computing a rooted phylogenetic network *N* that contains all input trees and minimizes the hybridization number. By construction, the network has the *tree-child* property which means that every internal node has at least one child for which it is the only parent. ALTS is a fast heuristic, rather than an exact algorithm, because it does not consider all possible orderings *π* of the input taxa, but rather uses a greedy heuristic to obtain a good ordering, and because it uses a progressive-alignment-like heuristic to perform the shortest-common superstring calculation, rather than solving this problem exactly.

The ALTS algorithm requires that all input trees are fully resolved. This poses a severe limitation in practice, because low-confidence edges in the input trees cannot be removed in a preprocessing step and then usually lead to unnecessarily complicated reticulation scenarios in the output. Also, the ALTS algorithm does not allow missing taxa in input trees.

Our new PhyloFusion algorithm accommodates input trees with multifurcations and unequal taxon sets. It is obtained by modifying the ALTS algorithm as follows:

0 Two additional preprocessing steps are undertaken.
  - All low-confidence input edges are contracted. By default, a threshold of 70% support is used.
  - All input trees are subjected to mutual refinement so as to resolve input nodes in a consistent manner.
1 There are two modifications of the first step:
  - The root node of tree *t* is labeled by the smallest taxon that is contained in the tree.
  - If *u* is an internal tree node, with children *v*_1_, *v*_2_, … *v*_*k*_ (*k ≥* 2), then define its label as

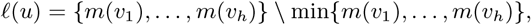

the set of all but the smallest taxon in {*m*(*v*_1_), …, *m*(*v*_*h*_)}.
2 The second step is modified as follows. The label *𝓁*(*v*) associated with an internal node is a set containing one or more taxa. Any taxon *x* contained in a given tree *t* occurs exactly twice, once in the label of some internal node *p*(*x*) and once in the label of a leaf *q*(*x*). Let *lth*(*x*) = (*𝓁*(*v*_1_), …, *𝓁*(*v*_*k*_)) denote the sequence of labels obtained from the interior nodes *v*_1_, …, *v*_*k*_ on the path from *p*(*x*) to *q*(*x*). We will refer to this as the “lineage taxon hypersequence” because the elements are sets rather than individual taxa. During the computation of a shortest common super sequence, two such sets may only be “aligned” to each other if they are in a containment relationship, that is if one contains the other.
3 Modification of the third step: For every taxon *x*, the elements of the list *sssc*(*x*) are sets of taxa.
4 Modification of the fourth step: For each taxon *x* in the order of *π* and for each internal node *v* in the path from *p*(*x*) to *q*(*x*), connect *v* to the first node *w* with label *𝓁*(*w*) = {*x*}, for all *x ∈ 𝓁*(*v*).

The goal is to compute a tree-child network that displays all the input trees. If one input tree contains a multifurcation and another input tree contains a resolution of the same node, then this will lead to multiple reticulations in the network that are necessary to accommodate both the multifurcation and its resolution. To reduce the number of reticulations that arise in this way, in the preprocessing step 0’ we mutually resolve the nodes of the input trees where possible.

To accommodate trees that have overlapping, but unequal taxon sets, the key point is that, in step 2’, we label the root by the smallest taxon contained in the current tree, rather than by the smallest taxon *x*_*π*(1)_. For any taxon that not in the current tree, its lineage taxon hypersequence is set to the empty sequence.

In practical applications, it is useful to have the edge weights in the network *N* reflect the average weights of the corresponding input edges. This can be solved using linear programming [Zhang et al., 2023]. In our implementation, we solve this by extracting, for each edge *e* in *N*, the set of all softwired clusters that it represents [Huson et al., 2012], and then averaging over the weights associated with each such cluster in the set of input trees. Because the number of softwired clusters contained in a rooted network grows exponentially with the number of reticulations, this calculation is only feasible for low numbers of reticulations.

In both ALTS and PhyloFusion, different orderings *π* of the taxa may give rise to different hybridization numbers. In PhyloFusion, we provide four heuristic modes, *fast, medium* or *thorough* (default, used in the analyses reported in this paper), using 10 *× n*, 150 *× n*, and 300 *× n* different random orderings, respectively, and evaluate these in parallel.

We have implemented the PhyloFusion algorithm in the SplitsTree app [Huson and Bryant, 2024]. SplitsTree allows the user to load a collection of phylogenetic trees, select a subset of trees, root them by outgroup or midpoint, contract edges using a confidence threshold, and then run the PhyloFusion algorithm to obtain a rooted phylogenetic network, which then can be displayed side-by-side with the input trees. Optionally, the algorithm can return multiple networks found during the heuristic search, all having the same hybridization number.

To create the input trees used for performance evaluation in Figure 2, we downloaded a multiple sequence alignment of 470 DNA sequences of length 829 from http://www.atgc-montpellier.fr/phyml/benchmarks/data/big_nucleic/ and then used IQ-TREE [Minh et al., 2020], default options, to compute a maximum-likelihood tree *T*, which we rooted at its midpoint. To obtain one input dataset for

PhyloFusion, for a given number of taxa *n*, we extracted the subtree *T*_*n*_ that is induced by the first *n* taxa of *T*. Then, for a given number *t* = 2, 5, 10, we created *t* new trees 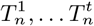, each one obtained from a copy of *T*_*n*_, by randomly removing *m* = 0 or *m* = 5% of its taxa, then randomly contracting *c* = 0 or *c* = 10% of its internal edges, and then finally, by applying a random rSPR modification [Bordewich and Semple, 2005] to it. The code that we developed to perform this study is available as a program called `sample-trees` that we make available as part of the SplitsTree package. The algorithm was run on a MacBook Pro laptop (M2 chip) using 8 cores.

Based on this construction, the minimum hybridization number of any rooted phylogenetic network *N* containing 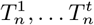 will be *n*, or less, in the rare case that some of the applied rSPRs undo each other. To address this, we used max(*n, h*) as hybridization number for a computed network to create the performance plots.

We also compared PhyloFusion against the Autumn algorithm [Huson and Linz, 2018] for two input trees, over the range of 10-200 taxa, based on the above mentioned tree with 470 taxa. Each pair of input trees differed by one rSPR per 10 taxa, and each tree was subjected to 10% missing taxa and 10% contracted internal edges. While the Autumn algorithm was able to compute an optimal network on datasets with up to 60 taxa and harboring at most 6 rSPRs, the algorithm timed out after 10 minutes on all but one of the larger datasets. In contrast, PhyloFusion was always able to run to completion on all datasets, requiring just over 200 seconds for 200 taxa, and achieving on average *h* = 27 where the number of rSPRs was 20.

## Contributions

L.Z. designed the algorithm. B.C. and D.H.H. implemented the algorithm in SplitsTree and performed data analysis. D.H.H. and L.Z. wrote the initial draft of the manuscript and all authors contributed to it.

## Acknowledgements

Research was partially completed while the authors were visiting the Institute for Mathematical Sciences, National University of Singapore in 2023. L.Z. was supported by The Singapore MOE Academic Research Fund Tier 1 [A-8001951-00-00].

## References

H.J. Bandelt, P. Forster, and A. Röhl. Median-joining networks for inferring intraspecific phylogenies. Molecular Biology and Evolution, 16:37–48, 1999.

Magnus Bordewich and Charles Semple. On the computational complexity of the rooted subtree prune and regraft distance. Annals of Combinatorics, 8(4):409–423, 2005.

D. Bryant and V. Moulton. Neighbor-net: An agglomerative method for the construction of phylogenetic networks. Molecular Biology and Evolution, 21(2):255–265, 2004.

M. Gruenstaeudl. Why the monophyly of nymphaeaceae currently remains indeterminate: an assessment based on gene-wise plastid phylogenomics. Plant Systematics and Evolution, 305:827–836, 2019.

D.H. Huson and D. Bryant. The SplitsTree app: interactive analysis and visualization with phylogenetic trees and networks. Under review, 2024.

D.H. Huson and S. Linz. Autumn algorithm—computation of hybridization networks for realistic phylogenetic trees. IEEE/ACM Transactions on Computational Biology and Bioinformatics, 15:398–420, 2018.

D.H. Huson, R. Rupp, and C. Scornavacca. Phylogenetic Networks. Cambridge, 2012.

S. Koblmueller, N. Duftner, K. M. Sefc, M. Aibara, M. Stipacek, M. Blanc, B. Egger, and C. Sturmbauer. Reticulate phylogeny of gastropod-shell-breeding cichlids from lake tanganyika-the result of repeated introgressive hybridization. BMC evolutionary biology, 7:7, 2007.

B. Q. Minh, H. A. Schmidt, O. Chernomor, D. Schrempf, M. D. Woodhams, A. von Haeseler, and R. Lanfear. IQ-TREE 2: New models and efficient methods for phylogenetic inference in the genomic era. Molecular Biology and Evolution, 37(5):1530–1534, 2020.

M. Stolzer, H. Lai, M. Xu, D. Sathaye, B. Vernot, and D. Durand. Inferring duplications, losses, transfers and incomplete lineage sorting with nonbinary species trees. Bioinformatics, 28(18):i409–i415, 2012.

Leo van Iersel, Remie Janssen, Mark Jones, Yukihiro Murakami, and Norbert Zeh. A practical fixed-parameter algorithm for constructing tree-child networks from multiple binary trees. Algorithmica, 84:917–960, 2022. doi: 10.1007/s00453-022-00959-0. URL https://doi.org/10.1007/s00453-022-00959-0.

L. Zhang, N. Abhari, C. Colijn, and Y. Wu. A fast and scalable method for inferring phylogenetic networks from trees by aligning lineage taxon strings. Genome Res, 2023.

